# Coordination and assembly of protein complexes encoded across mitochondrial and nuclear genomes is assisted by CLPP2 in *Arabidopsis thaliana*

**DOI:** 10.1101/2020.01.22.907055

**Authors:** Jakob Petereit, Owen Duncan, Monika W Murcha, Ricarda Fenske, Emilia Cincu, Jonathan Cahn, Adriana Pružinská, Aneta Ivanova, Laxmikanth Kollipara, Stefanie Wortelkamp, Albert Sickmann, Jiwon Lee, Ryan Lister, A Harvey Millar, Shaobai Huang

**Affiliations:** ARC Centre of Excellence in Plant Energy Biology, School of Molecular Sciences, The University of Western Australia, 35 Stirling Highway, WA, 6009, Australia; Leibniz-Institut für Analytische Wissenschaften – ISAS – e.V., Dortmund, Germany; Department of Chemistry, College of Physical Sciences, University of Aberdeen, Aberdeen, Scotland, United Kingdom; Medizinische Fakultät, Medizinische Proteom-Center (MPC), Ruhr-Universität Bochum, Bochum, Germany; Centre for advanced Microscopy, The Australian National University, Acton, ACT 2601, Australia; The Harry Perkins Institute of Medical Research, Perth, WA 6009, Australia

## Abstract

Protein homeostasis in eukaryotic organelles and their progenitor prokaryotes is regulated by a series of proteases including the caseinolytic protease (CLPP). CLPP has essential roles in chloroplast biogenesis and maintenance, but the significance of the plant mitochondrial CLPP remains unknown and factors that aid coordination of nuclear and mitochondrial encoded subunits for complex assembly in mitochondria await discovery. We generated knock-out lines of the single gene for the mitochondrial CLP protease subunit, *CLPP2,* in *Arabidopsis thaliana*. Mutants had higher abundance of transcripts from mitochondrial genes encoding OXPHOS protein complexes, while transcripts for nuclear genes encoding other subunits of the same complexes showed no change in abundance. In contrast, the protein abundance of specific nuclear-encoded subunits in OXPHOS complexes I and V increased in CLPP2 knockouts, without accumulation of mitochondrial-encoded counterparts in the same complex. Protein complexes mainly or entirely encoded in the nucleus were unaffected. Analysis of protein import, assembly and function of Complex I revealed that while function was retained, protein homeostasis was disrupted through decreased assembly, leading to accumulation of soluble subcomplexes of nuclear-encoded subunits. Therefore, CLPP2 contributes to the mitochondrial protein degradation network through supporting coordination and assembly of protein complexes encoded across mitochondrial and nuclear genomes.

**One sentence summary:** CLPP contributes to the mitochondrial protein degradation network through supporting coordination and assembly of protein complexes encoded across mitochondrial and nuclear genomes.

## Introduction

Mitochondria contain numerous protein complexes that catalyse important functions including oxidative phosphorylation leading to ATP synthesis, respiratory carbon metabolism, consumption of oxygen and the production of reactive oxygen species (ROS) (Millar et al., 2011; Huang et al., 2016). Subunits for some of these protein complexes are encoded across mitochondrial and nuclear genomes. This requires separate gene expression, protein synthesis and protein import into mitochondria followed by a coordinated complex assembly. The regulation at a transcriptional level is limited, owing to the general lack of mitochondrial gene specific transcript activation (Costanzo and Fox, 1990; Giegé et al., 2000; Kühn et al., 2009).

Some of the coordination of nuclear and mitochondrial encoded protein complex assembly occurs at the translation level and is unidirectional (Couvillion et al., 2016), with cytosolic events influencing mitochondrial mRNA levels. In Arabidopsis cell culture under carbon starvation and refeeding conditions, a transient dysregulation in the stoichiometry of Complex V subunits ATP1 and ATP2, encoded by mitochondria and nuclear genomes respectively, has been identified (Giege et al., 2005). *ATP1* was increased at a transcript level but not at the level of ATP1 protein abundance, while ATP2 increased at the protein level, but the *ATP2* transcript was unchanged. Giegé et al (2005) proposed that post-translational processes in assembly were responsible for rebalancing this process after re-feeding of cells but did not identify factors responsible for it. It has previously been suggested that unassembled protein complex subunit degradation by mitochondrial proteolysis is the basis for such regulation (Sarria et al., 1998), but the specific protease(s) involved have not been identified.

Organellar protease networks exist in mitochondria and plastids to control intra-organellar protein degradation. (van Wijk, 2015) ATP dependent AAA+ (ATPase-associated with various cellular activities) endopeptidases such as LON1 (long-filament phenotype-1), CLPP (caseinolytic protease) and FTSH (filamentous temperature sensitive H) are the dominant components of these networks that are conserved across eukaryotes. In plants, mitochondrial LON1 acts as a chaperone aiding the proper folding of newly synthesized/imported proteins by stabilising them and as a protease by degrading mitochondrial protein aggregates (Li et al 2017). Loss of mitochondrial FTSHs leads to oxidative stress, and FTSH are shown to be components of the defence against accumulation of carbonylated proteins in plant mitochondria (Smakowska et al., 2016). To date, there have not been any reports of the effect of loss of CLPP on plant mitochondria.

The CLPP protease has an active Ser-His-Asp catalytic triad and is widely present in bacteria, fungi, mammals and plants (Yu and Houry, 2007; Bhandari et al., 2018). The X-ray structure of CLPP has been solved from several different organisms with similar features: the protease subunit is comprised of two heptameric rings forming a cylinder-like structure which encloses a large chamber containing the protease active sites (Yu and Houry, 2007; Bhandari et al., 2018). Some bacteria such as the cyanobacterium *Synechococcus elongatus* contain CLPR which is an inactive version of CLPP lacking the catalytic triad (Schelin et al., 2002). In bacteria, CLPP forms complexes with AAA+ chaperones, CLPX and CLPA acts as a selective filter for specific target substrates (Gerth et al., 2008).

Plants have a diversified and complex CLP family with 6 active CLPP (CLPP1-6) and 4 inactive CLPR paralogs (CLPR1-4) found in *Arabidopsis thaliana* (Sjögren et al., 2006; van Wijk, 2015). In addition, Arabidopsis have CLP/HSP100 AAA+ chaperones with 7 class I (CLPB1-4, CLPC1-2, CLPD) in chloroplasts and 3 class II (CLPX1-3) in mitochondria (Peltier et al., 2004). Arabidopsis plastids have 5 CLPP proteases (CLPP1 encoded by the plastid genome and CLPP3-6) and 4 inactive CLPR, which together form the tetradecameric and asymmetric ∼350 kDa CLP protease core (van Wijk, 2015). Three CLP AAA+ chaperones (CLPC1-2, CLPD) and the adaptor CLPS1 deliver protein substrates to the protease complex for degradation (van Wijk, 2015). Plastid *clpp* and *clpr* mutants in Arabidopsis display severe phenotypes including disorder of embryogenesis, seedling development and chloroplast biogenesis. Complete loss of CLPP5 and CLPP4 is embryo lethal and the *clpp3* null mutant can only germinate and develop seedlings under heterotrophic conditions (Kim et al., 2013). CLPP1 is known to be essential for leaf development in tobacco (Shikanai et al., 2001). Knockout lines of either CLPR2 or CLPR4 cause embryogenesis delay and developmental arrest at the cotyledon stage (Kim et al., 2009). Upregulation of chaperone proteins and downregulation of photosynthesis coupled with upregulation of the ATP import pathway occur in *clpr2, clpr4 and clpr3* mutants (Kim et al., 2013). Dysfunction of CLPP6 in rice caused developmental stage-dependent virescent yellow leaf with decreased chlorophyll accumulation and impaired photosynthesis (Dong et al., 2013). Therefore, the plant CLPPR protease complex appears to play an essential role in plastid and chloroplast biogenesis and development.

Arabidopsis mitochondria contain a homotetradecameric CLPP2 core protease encoded by a single nuclear gene and three CLPX chaperones (CLPX1-3) (Peltier et al., 2004). The CLPP2 core has been detected by Blue native PAGE with a molecular mass of ∼320 kDa (Senkler et al., 2017), which is consistent with a predicted tetradecamer structure (Peltier et al., 2004), but no T-DNA lines for CLPP2 exist. In this study, we developed two knock-out lines of *CLPP2* in *Arabidopsis thaliana* by mutations using CRISPR-Cas9 technology, providing a tool for functional analysis of mitochondrial CLPP2. Our assessments using a range of omic technologies and specific follow up experiments have revealed that while CLPP2 is not essential for plant growth, it does contribute to the regulation of nuclear and mitochondrial protein complex assembly and maintenance in a manner consistent with the previously proposed post-translational processes that have been implicated in this control.

## Results

### CRISPR-Cas9 guided *CLPP2* knockout results in two stable knock-out mutants

The serine-type, ATP-dependent CLP protease system in Arabidopsis mitochondria contains a CLPP2 core subunit (AT5G23140) and three homologous CLPX chaperone subunits (CLPX1, AT5Gg53350; CLPX2, AT5G49840, CLPX3, AT1G33360) (van Wijk, 2015). So far, there is no reported or available T-DNA insertion line within the coding region for *CLPP2* (AT5G23140). In order to acquire independent stable *CLPP2* mutants, we used a CRISPR-Cas9 guided system to knock-out the CLPP2 core subunit (**Supplemental Figure 1**), resulting in two individual CRISPR-Cas9 mutants, *clpp2-1* and *clpp2-2*, showing a complete knock-out of the target gene. Both mutations occurred within exon 1 as shown by guides, primers and restriction sites/enzymes (Figure 1A).

**Figure 1:**
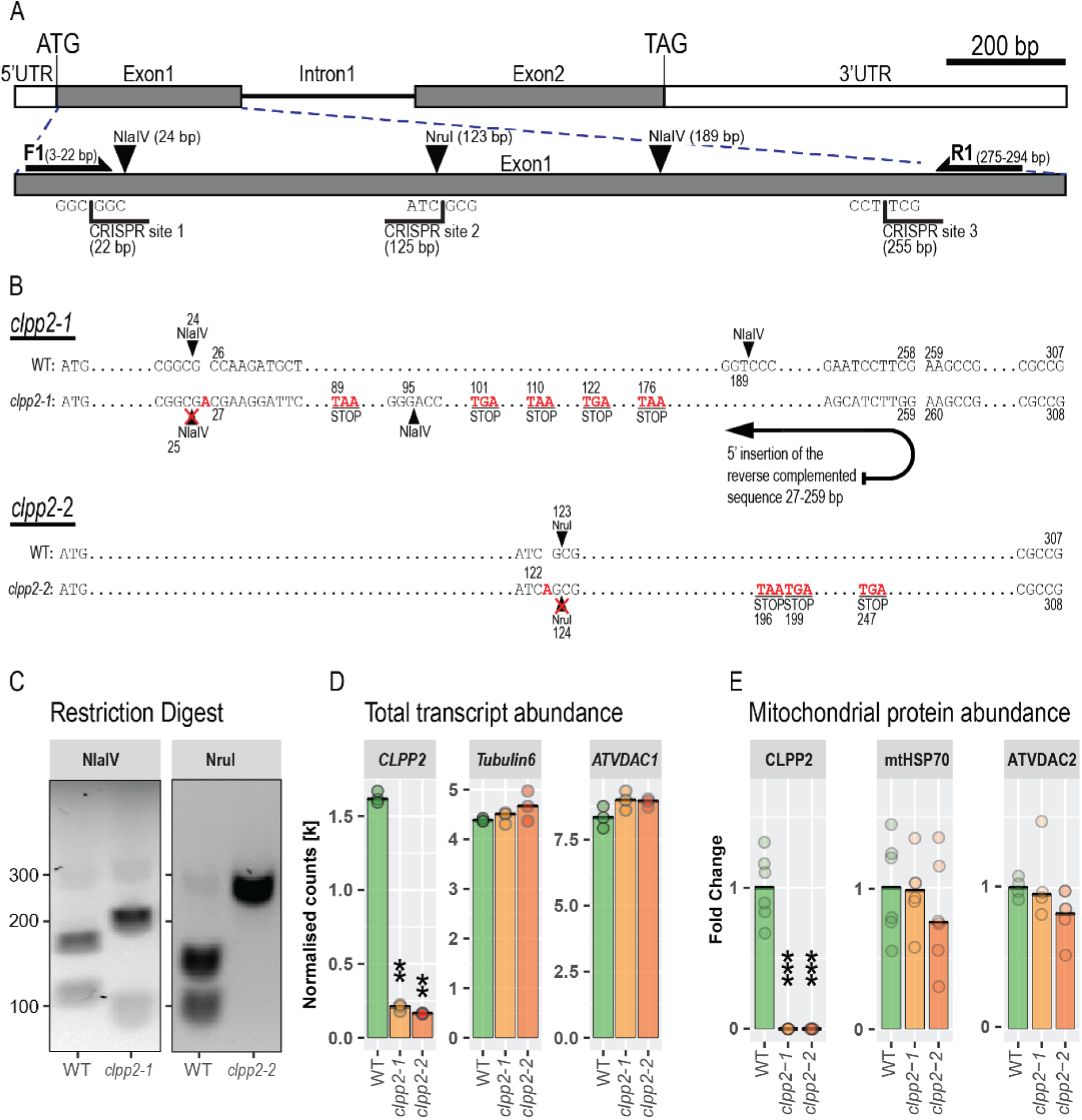
Development of two independent *CLPP2* mutants using CRISPR-Cas9. (**A**) The figure illustrates the layout of the CLPP2 gene structure with enlarged exon 1, showing CRISPR-Cas9target sites. Left/right arrows represent primer positions (F1 & R1) and downward arrows represent restriction sites used for restriction digest-based mutant confirmation of the mutations. (**B**) Mapping of CRISPR-Cas9 gene disruption for the two individual knock outs *clpp2-1* and *clpp2-2* (bottom). *Clpp2-1*: WT shows the exon 1 sequence map with the location of two NlaIV restriction sites. *Clpp2-1* has an adenine insertion (A) at CRISPR site 1 disrupting (X) one NlaIV restriction site and an insertion of the reverse complemented wild type sequence between CRISPR site 1 and CRISPR site 3 introducing 5 stop codons (TAA, TGA) resulting in change of the second NlaIV restriction site. *Clpp2-2*: WT shows the exon 1 sequence map with the location of one NruI restriction site. *Clpp2-2* has an adenine insertion (CRISPR site 2) resulting in the disruption of the NruI restriction site and insertion of 3 stop codons. (**C**) Confirmation of CRISPR-Cas9 gene disruption based on PCR fragment restriction digest amplified by F1 and R1 primers. NlaIV and Nrul digestion are shown in left and right, respectively. (**D**) Confirmation of CRISPR-Cas9 gene disruption at the transcript level based on RNA seq analysis. The normalised counts of the target gene *CLPP2*, and the two control genes *Tubulin6* and *ATVDAC1* are presented. ** represent p ≤ 0.01 (n = 3). (**E**) Confirmation of CRISPR-Cas9 gene disruption at the protein level based on a targeted proteomic approach. Fold changes of the target protein CLPP2, and the two control proteins mtHSP70 and ATVDAC2 are presented. *** represent p ≤ 0.0001 (n = CLPP2 : 3 peptides, 2 replicates, mtHSP70 : 3 peptides, 2 replicates, ATVDAC2 : 2 peptides, 2 replicates).

The first mutant *clpp2-1* showed two separate genetic events that caused the CLPP2 knock-out: firstly, a single adenine insertion at position of 25bp, introduced a frame shift and disrupted the restriction site of NlaIV at 24bp (Figure 1B); secondly, the introduction of the adenine at CRISPR site 1 created an identical 3’ end as the 5’end at CRISPR site 3. It is likely that this caused a second error from the nonhomologous end-joining (NHEJ) DNA repair mechanism or from the alternative nonhomologous end-joining DNA repair mechanism (MMEJ) known for insertions, deletions and inversions (McVey and Lee, 2008), inserting the complete fragment between CRISPR site 1 and CRISPR site 3 in a reverse complemented orientation. Consequentially, five downstream stop codons (TGA or TAA) were generated and the second NlaIV restriction site was shifted to the position at 95bp (Figure 1B). The second mutant *clpp2-2* had a single NHEJ error at CRISPR site 2 with an adenine insertion at the position of 124bp, resulting in a disruption of the NruI restriction site and the introduction of 3 downstream stop codons (TAA or TGA) (Figure 1B).

We confirmed the disruption of *CLPP2* in the mutants in the genetic sequence as well as its consequence on mRNA and protein levels. As the two mutations resulted in disruption of restriction sites within exon 1, mutants and WT genomic DNA was amplified using primers F1 and R1 (Figure 1A) and digested with NlaIV for *clpp2-1* and NruI for *clpp2-2*, respectively. NlaIV digested WT PCR product resulted in three fragments, a long 165bp fragment between the first and second NlaIV restriction site, an intermediate 105bp fragment from restriction site 2 to the 3’ end of PCR product and a short 24bp fragment (out of gel range) from the 5’ end to the restriction site 1 (Figure 1C). For *clpp2-1*, NIaIV restriction site 1 was disrupted and restriction site 2 moved in 5’ end direction. Accordingly, the restriction digest pattern had 2 fragments in total, a 199bp fragment from the new restriction site 2 to the 3’ end of the PCR product and a 92bp fragment from the 5’ end to the new restriction site 2 (Figure 1C). NruI digested WT exon 1 PCR product resulting in 2 fragments with 171bp and 120bp in length, while *clpp2-2* had a disrupted NruI restriction site and only showed the full 291bp PCR product (Figure 1C).

Based on RNAseq analysis, the transcript levels of *CLPP2* were significantly reduced in *clpp2-1* and *clpp2-2* to about 15% of the WT level (Figure 1D). The expression level of representative house-keeping genes such as general *TUBULIN6* (AT5G12250) and mitochondrial *ATVDAC1* (AT3G01280) were unaffected in both mutants (Figure 1D). While CRISPR-Cas9 systems shouldn’t actively alter transcript levels, the introduced frameshifts forming several stop codons into the sequence of *CLPP2,* more than 50bp away from the exon-exon junction (Figure 1B), created a high confidence position for exon-exon junction nonsense-mediated mRNA decay which may contribute to the observed low transcript abundance (Lloyd, 2018).

To detect the protein abundance of CLPP2, we isolated mitochondria and conducted multiple reaction monitoring (MRM)-based targeted proteomic analysis and included the theoretical peptide of the inverted CLPP2-1 section. The peptide MRM transitions were presented in **Supplemental Table 1**. CLPP2 protein abundance in mitochondria from both mutants was undetectable, while all selected peptides of CLPP2 were detected in WT (Figure 1E). We also did not detect any evidence of theoretical CLPP2-1 peptides. Two representative mitochondrial housekeeping proteins (mtHSP70-AT4G37910: mitochondrial heat shock protein 70; ATVDAC2-AT5G67500: voltage dependent ion channel) showed no significant change in protein abundance in either mutant (Figure 1E). The virtual absence of CLPP2 proteins in both mutants could be explained by the introduction of stop codons at the first exon (Figure 1B) that fully disrupted the translation of any functional CLPP2 protein.

### Loss of CLPP2 has an impact on transcript levels of mitochondrial genes but little effect on expression of nuclear genes

To evaluate global transcript abundance changes upon the loss of the mitochondrial CLPP2, we used high-throughput RNA-sequencing to find and quantify differentially expressed genes (DEGs). We analysed the transcriptome of hydroponically grown seedlings of *clpp2-1*, *clpp2-2* and WT, with three biological replicates for each genotype. Based on a filter of a log2 fold change (log2FC) exceeding ±0.4 and adjusted p-values ≤ 0.05, we identify 62 DEGs using the R package Deseq2 (Love et al., 2014), which showed a consistent pattern in both mutants (Figure 2). Only 0.1% of nuclear gene transcripts had significant changes in abundance in both *clpp2-1 and clpp2-2*. This was considerably lower than the 33% of mitochondrial transcripts that showed significant changes in abundance in both mutants (Figure 2), indicating a specific impact of the disruption of *CLPP2* on the abundance of transcripts from mitochondrial genes.

**Figure 2:**
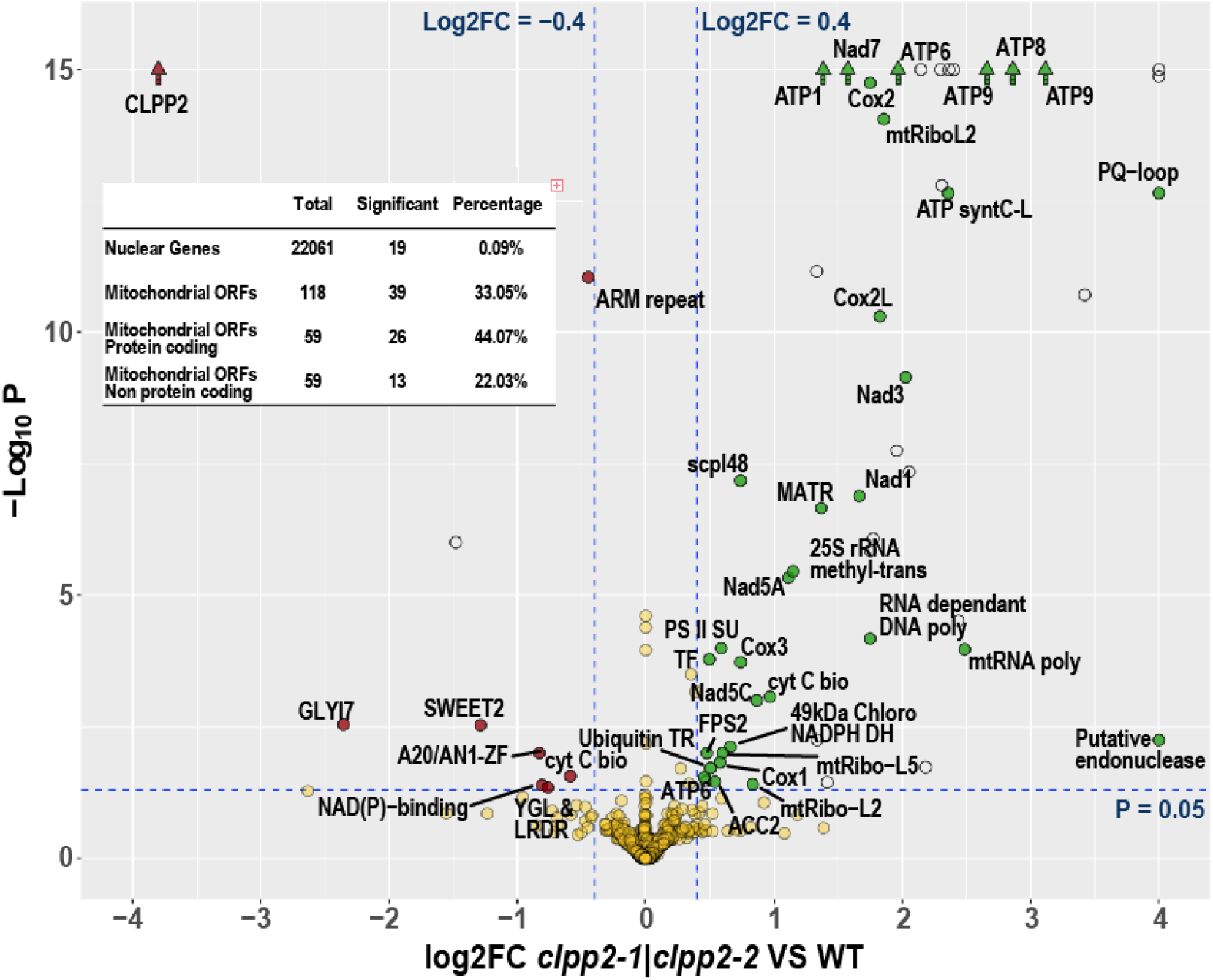
Changes in global transcripts abundances in *CLPP2* mutants compared with WT. The volcano plot illustrates overlapping significant changes of total transcript abundances of the two KO-mutants compared to the wild type. The negative Log10 transformed adjusted p-values (p.adj) are plotted against the log2 fold change (log2FC) of the transcript abundances measured by RNAseq. Dashed lines separate gene transcripts with significant changes in abundance with the log2FC cut off of ±0.4 and adjusted p-value cut off of 0.05. Depleted gene transcripts are shown in red and accumulated gene transcripts in green. Transcripts with p.adj values under 1×10^-15^ are displayed as arrow shapes. Transposable elements and hypothetical proteins are displayed as empty circles. The inserted table displays the summary total number of detected nuclear and mitochondrial genes and the overlapped numbers with significant changes in abundance from both mutants against the wild type.

In *clpp2-1 and clpp2-2*, 41 of the 52 upregulated DEGs from nuclear and mitochondrial genomes encode mitochondrial-localised proteins, while 3 of 8 downregulated DEGs, other than CLPP2, encodes a mitochondrial-localised protein (**Table 1**). Two genes that encode non-mitochondrial targeted proteins and have unknown function stand out, showing very high and consistent log2FC values. AT4G36850 encodes a conserved PQ-Motif protein (Pfam: PF04193, involved in lysosome targeting) protein and AT3G56730 encodes a putative endonuclease with a conserved LabA-like PIN domain of limkain b1 (cd10910), which is a common target of human antibodies targeting cytoplasmic vesicle-like structures (Marchler-Bauer et al., 2017; El-Gebali et al., 2018).

**Table 1:** Functional category of genes with significant changes in transcript abundances exceeding the cut off in clpp2-1 and clpp2-2. Genes were classified into the following groups: OXPHOS (mitochondrial oxidative phosphorylation components); mtBIO (mitochondrial biogenesis); KO (nuclear encoded CLPP2 protease knocked out (KO) in the mutants); other (ungrouped genes). The log2FC for both mutants compared to the wild type and the corresponding adjusted p-values are listed. Arabidopsis gene identifiers (AGI), short descriptions and the subcellular localisation of the protein based on the SUBA4 database are shown (M: Mitochondrion, N: Nucleus, Cyt: Cytosol, P: Plastid, PM: Plasma Membrane, EX: Extracellular, V: Vacuole, ER: Endoplasmic Reticulum).

| Group | Symbol | AGI | Description | Location | log2FC<br>*clpp2-1* | log2FC<br>*clpp2-2* | p.adj<br>*clpp2-1* | p.adj<br>*clpp2-2* |
| --- | --- | --- | --- | --- | --- | --- | --- | --- |
| OXPHOS | ATP9 | ATMG01090 | ATP synthase 9 | M | 3.1 | 3.2 | 0.00 | 0.00 |
| OXPHOS | ATP8 | ATMG00480 | Atp8 | M | 3.1 | 2.9 | 0.00 | 0.00 |
| OXPHOS | ATP9 | ATMG01080 | F0-ATPase subunit 9 | M | 2.8 | 2.7 | 0.00 | 0.00 |
| OXPHOS | ATPSyntC | ATMG00040 | ATP synthase subunit C | M | 2.4 | 3.0 | 0.00 | 0.00 |
| OXPHOS | Nad3 | ATMG00990 | NADH dehydrogenase 3 | M | 2.0 | 2.0 | 0.00 | 0.00 |
| OXPHOS | ATP6 | ATMG00410 | ATPase subunit 6-1 | M | 2.0 | 2.1 | 0.00 | 0.00 |
| OXPHOS | Cox2L | ATMG01280 | Cytochrome C oxidase subunit II-like | M | 1.8 | 2.3 | 0.00 | 0.00 |
| OXPHOS | Cox2 | ATMG00160 | cytochrome oxidase 2 | M | 1.7 | 2.0 | 0.00 | 0.00 |
| OXPHOS | Nad1 | ATMG00516 | NADH dehydrogenase 1C | M | 1.7 | 1.7 | 0.00 | 0.00 |
| OXPHOS | Nad7 | ATMG00510 | NADH dehydrogenase subunit 7 | M | 1.6 | 2.1 | 0.00 | 0.00 |
| OXPHOS | ATP1 | ATMG01190 | ATP synthase subunit 1 | M | 1.4 | 1.8 | 0.00 | 0.00 |
| OXPHOS | Nad5A | ATMG00513 | NADH dehydrogenase 5A | M | 1.1 | 1.5 | 0.00 | 0.00 |
| OXPHOS | cyt C bio | ATMG00180 | cytochrome C biogenesis 452 | M | 1.0 | 1.4 | 0.00 | 0.00 |
| OXPHOS | Nad5C | ATMG00060 | NADH dehydrogenase subunit 5C | M | 0.9 | 1.3 | 0.00 | 0.00 |
| OXPHOS | Cox3 | ATMG00730 | cytochrome c oxidase subunit 3 | M | 0.7 | 1.2 | 0.00 | 0.00 |
| OXPHOS | Cox1 | ATMG01360 | cytochrome oxidase | M | 0.6 | 0.9 | 0.02 | 0.00 |
| OXPHOS | ATP6 | ATMG01170 | ATPase | M | 0.5 | 0.9 | 0.03 | 0.00 |
| OXPHOS | cyt C bio | ATMG00900 | cytochrome C biogenesis 256 | M | -1.4 | -0.6 | 0.00 | 0.03 |
| mtBIO | mtRNA poly | ATMG00490 | Mitovirus RNA-dependent RNA polymerase | M | 2.5 | 2.9 | 0.00 | 0.00 |
| mtBIO | mtRibo L2 | ATMG00980 | Ribosomal protein S12/S23 | M | 1.9 | 1.9 | 0.00 | 0.00 |
| mtBIO | DNA/RNA polymerase | ATMG00810 | DNA/RNA polymerases | M | 1.7 | 2.2 | 0.00 | 0.00 |
| mtBIO | 25S rRNA methyl-trans | AT4G26600 | methyltransferases | N | 1.5 | 1.1 | 0.00 | 0.00 |
| mtBIO | MATR | ATMG00520 | Intron maturase | M | 1.4 | 1.6 | 0.00 | 0.00 |
| mtBIO | mtRibo L2 | ATMG00560 | Nucleic acid-binding | M | 0.8 | 1.5 | 0.04 | 0.00 |
| mtBIO | mtRibo L5 | ATMG00210 | ribosomal protein L5 | M | 0.6 | 1.0 | 0.01 | 0.00 |
| other | PQ-Motif | AT4G36850 | PQ-loop repeat family protein | PM | 16.0 | 16.9 | 0.00 | 0.00 |
| other | endonuclease | AT3G56730 | endonuclease or glycosyl hydrolase | N,Cyt | 6.7 | 8.1 | 0.01 | 0.00 |
| other | scpl48 | AT3G45010 | serine carboxypeptidase-like 48 | EX | 1.2 | 0.7 | 0.00 | 0.00 |
| other | 49kDa Chl NADPH DH | ATCG01110 | 49 kDa SU Chloroplast NAD(P)H-DH | P | 0.7 | 0.9 | 0.01 | 0.00 |
| other | ACC2 | AT1G36180 | acetyl-CoA carboxylase 2 | P | 0.6 | 0.5 | 0.03 | 0.01 |
| other | PSII SU | ATCG00280 | photosystem II reaction center protein C | P | 0.6 | 0.9 | 0.00 | 0.00 |
| other | FPS2 | AT4G17190 | farnesyl diphosphate synthase 2 | M,Cyt | 0.5 | 0.5 | 0.01 | 0.00 |
| other | Ubiquitin TR | AT5G63760 | RING/U-box superfamily protein | N | 0.5 | 1.0 | 0.02 | 0.00 |
| other | TF | AT5G27410 | D-aminoacid aminotransferase-like | Cyt | 0.5 | 0.6 | 0.00 | 0.00 |
| other | NAD(P)-binding | AT2G17845 | NAD(P)-binding Rossmann-fold | M | -0.8 | -2.0 | 0.04 | 0.00 |
| other | ARM repeat | AT3G56210 | ARM repeat superfamily protein | M | -0.8 | -0.4 | 0.00 | 0.00 |
| other | A20/AN1-ZF | AT1G12440 | A20/AN1-like zinc finger family protein | N | -1.1 | -0.8 | 0.01 | 0.01 |
| other | SWEET2 | AT3G14770 | Nodulin MtN3 family protein | V | -1.3 | -1.3 | 0.00 | 0.00 |
| other | YGL&LRDR | AT3G15450 | YGL and LRDR motifs | Per,Cyt | -1.9 | -0.8 | 0.00 | 0.05 |
| other | GLYI7 | AT1G80160 | Lactoylglutathione lyase | Cyt,ER,PM | -2.4 | -2.8 | 0.00 | 0.00 |
| KO | CLPP2 | AT5G23140 | CLPP2 | M | -4.3 | -4.7 | 0.00 | 0.00 |

Out of the 41 upregulated mitochondrial-localised DEGs, 24 originate from the mitochondrial genome (**Table 1**) and encode proteins of the OXPHOS system, mitochondrial biogenesis apparatus and various hypothetical proteins, transposons and retro-transposons. Notably, mitochondrial DEGs encoding Complex V ATP synthase subunits (such as ATP1, ATP6 (AT3G46430, AT5G59613), ATP8, ATP9) showed higher expression levels in both *CLPP2* mutants compared to WT (**Table 1**). Some mitochondrial genes encoding Complex I subunits (NAD1, 3, 5, 7) and Complex IV subunits (COX1, 2, 3) were also observed with high expression level in both mutants (**Table 1**). Some mitochondrial DEGs were annotated with RNA or DNA polymerase (ATMG00490, and ATMG00810) and intron maturase (ATMG00520) functions (**Table 1**).

### Loss of CLPP2 alters mitochondrial protein homeostasis

Changes in transcript abundance often show only a poor correlation with the abundance of finally accumulated proteins (Haider and Pal, 2013). To identify differentially expressed proteins (DEPs) in isolated mitochondria from hydroponically grown Arabidopsis seedlings, we conducted a quantitative proteomic approach. We detected 797 proteins from mitochondrial membrane fractions in mutants and WT (**See Supplemental Table 2**). Among them, we identified 22 DEPs with increased abundance and 5 DEPs with decreased abundance. These showed a consistent pattern in *clpp2-1* and *clpp2-2* with a log2 fold change (log2FC) exceeding ±0.4 and p-values ≤ 0.05 (Figure 3). Details of the 27 DEPs are listed in **Table 2** and are grouped based on functional categories. We detected the accumulation of ATP2 (AT5G08690) for Complex V and four subunits of Complex I (24kDa: AT4G02580, 51kDa: AT5G08530, 75kDa: AT5G37510 and B14: AT3G12260) (**Table 2**). Three of the over-accumulated Complex I subunits belong to the N-module of the matrix arm, which is assembled before it is attached to the membrane arm of Complex I (Ligas et al., 2019). We observed an accumulation of proteins related to protein synthesis, such as three components of the mito-ribosome (AT4G3090, AT1G61870 and AT5G64670) and the putative RNA helicase AT3G22130 (RH9) as well as mitochondrial proteases, such as MPPα-1,α-2; MPPβ, CLPX-1 and PREP1 (**Table 2**). MPPs are mitochondrial peptidases that are embedded in Complex III and are responsible for mitochondrial presequence cleavage after protein import (Braun et al., 1992; Zhang et al., 2001). CLPX-1 is the chaperone subunit of the CLPXP complex (van Wijk, 2015) and PREP1 degrades mitochondrial pre-sequences after their cleavage from proteins (Kmiec and Glaser, 2012; Kmiec et al., 2014). A change to the abundance of these proteases in the absence of CLPP2 could influence the degradative landscape within the mitochondria of the *CLPP2* mutants. An analysis of matched samples of the soluble mitochondrial proteome produced similar results (including changes in the abundance of 24 kDa subunit, RH9, CLPX1, mtHSP70-1 and PPR proteins), albeit with different ratios of abundance for a range of proteins (including MPP, ATP2 and ATP3 (**Supplemental Figure 2, Supplemental Table 2).**

**Figure 3:**
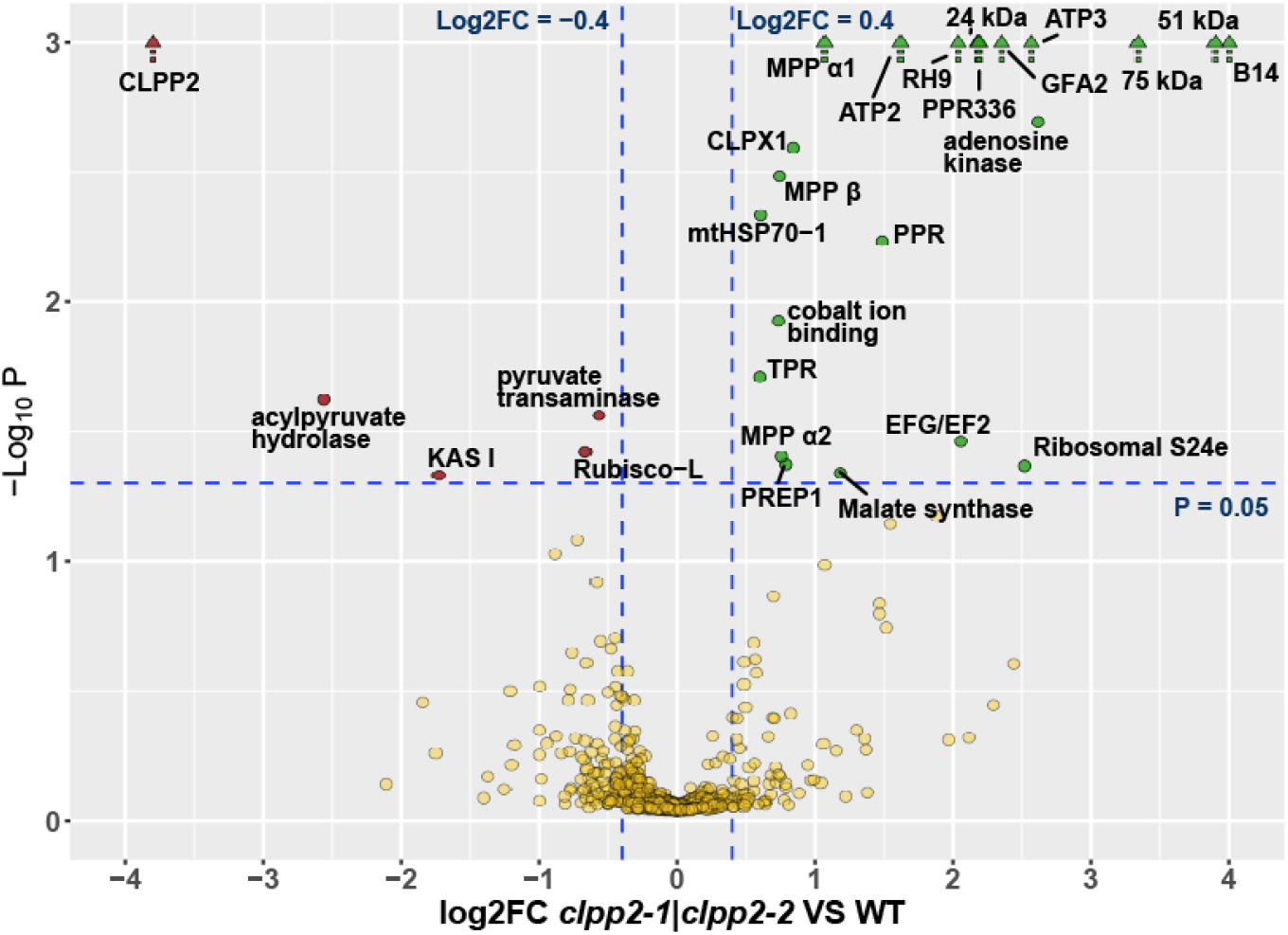
Changes in protein abundances in membrane fraction of isolated mitochondria in *CLPP2* mutants compared with WT. The volcano plot illustrates overlapping significant changes of LFQ protein intensities of the two KO-mutants compared to the wild type, using a log2 fold change (log2FC) cut off of ±0.4 and an adjusted p-value cut off of 0.05 (dashed blue lines). Depleted proteins are shown in red and accumulated proteins in green. Proteins with adjusted p-values under 1×10^-15^ are displayed as arrow shapes.

**Table 2:**
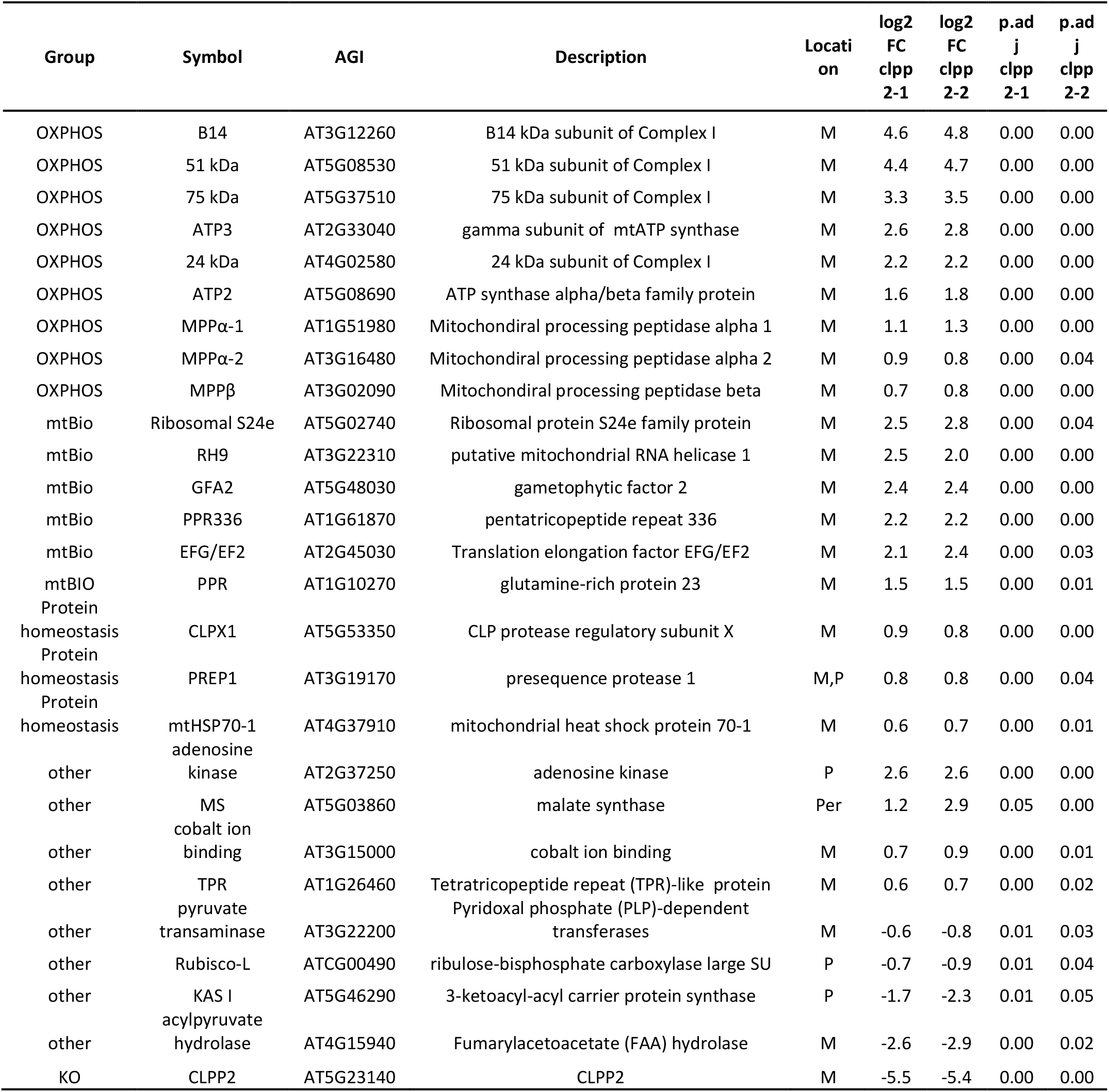
Functional category of proteins with significant changes in abundance in clpp2-1 and clpp2-2. Proteins were classified into the following groups: OXPHOS (mitochondrial oxidative phosphorylation components); mtBIO (mitochondrial biogenesis); Protein homeostasis; KO (nuclear encoded CLPP2 protease knocked out (KO) in the mutants); other (ungrouped genes). The log2 fold changes in protein abundance in both mutants compared to wild type and the corresponding adjusted p-values are listed. Arabidopsis gene identifiers (AGI), short descriptions and the subcellular protein localisation based on the SUBA4 database (M: Mitochondrion, N: Nucleus, P: Plastid, Cyt: Cytosol, PM: Plasma Membrane, EX: Extracellular, V: Vacuole, ER: Endoplasmic Reticulum) are presented.

To investigate a potential modification in pre-sequence processing or partial degradation of mature proteins, we conducted a combined approach of protein labelling with isobaric tags for relative and absolute quantification (Ross et al., 2004) and charge based fractional diagonal chromatography (ChaFRADIC) (Venne et al., 2013; Venne et al., 2015) to analyse the N-terminal mature sequence of mitochondrial proteins. Sequence logo analysis indicated that there was no difference in pre-sequences cleavage as there were very conserved −2R and −3R cutting sites detected in mitochondrial proteins in both mutants and WT (**Supplemental Figure 3A & 3B)**. There was also no apparent difference in the abundance of any pre-sequence peptides between mutants and WT (**Supplemental Figure 3C**). In addition, no significant difference was observed in cleavage sites in the middle or C-terminus of protein sequence between the two mutants and WT (**Supplemental Figure 3C & 3D**). Therefore, loss of CLPP2 had no measurable impact on mitochondrial protein maturation and left no partially degraded parts of mitochondrial proteins that we could detect.

### Loss of CLPP2 disrupts the coordination of mitochondria- and nuclear-encoded subunits of respiratory and ribosomal complexes

To directly compare between mitochondrial transcript and protein abundance changes for individual subunits located in the same protein complex, we compiled the changes in transcript level and protein abundance for all subunits of the mitochondrial OXPHOS system and the mitochondrial ribosome (Figure 4). Complex I, the largest protein complex of the mitochondrial OXPHOS system, is encoded by nine mitochondrial encoded genes and 47 nuclear encoded genes (Ligas et al., 2019; Meyer et al., 2019). Our data showed a consistent transcript upregulation for five mitochondrial encoded subunits (NAD1, 3, 5, 7) in both mutants, but we did not find NAD7 protein accumulation in either mutant (Figure 4A). In contrast, complex I subunits (24 kDa, 51 kDa, 75 kDa and B14), encoded by four nuclear genes, were highly accumulated in protein abundance with no change in transcript levels (Figure 4A). A similar pattern was observed for Complex V; five (ATP1, ATP6 (AT3G46430, AT5G59613), ATP7, ATP8) out of six mitochondrial transcripts had higher abundance in both mutants, but ATP1 and ATP8 did not show any accumulation at the protein level (Figure 4E). There was no consistent transcript upregulation of nuclear genes encoding Complex V subunits in both mutants, but there was high accumulation of the ATP2 protein, encoded by the nuclear genome, in both mutants (Figure 4E).

**Figure 4:**
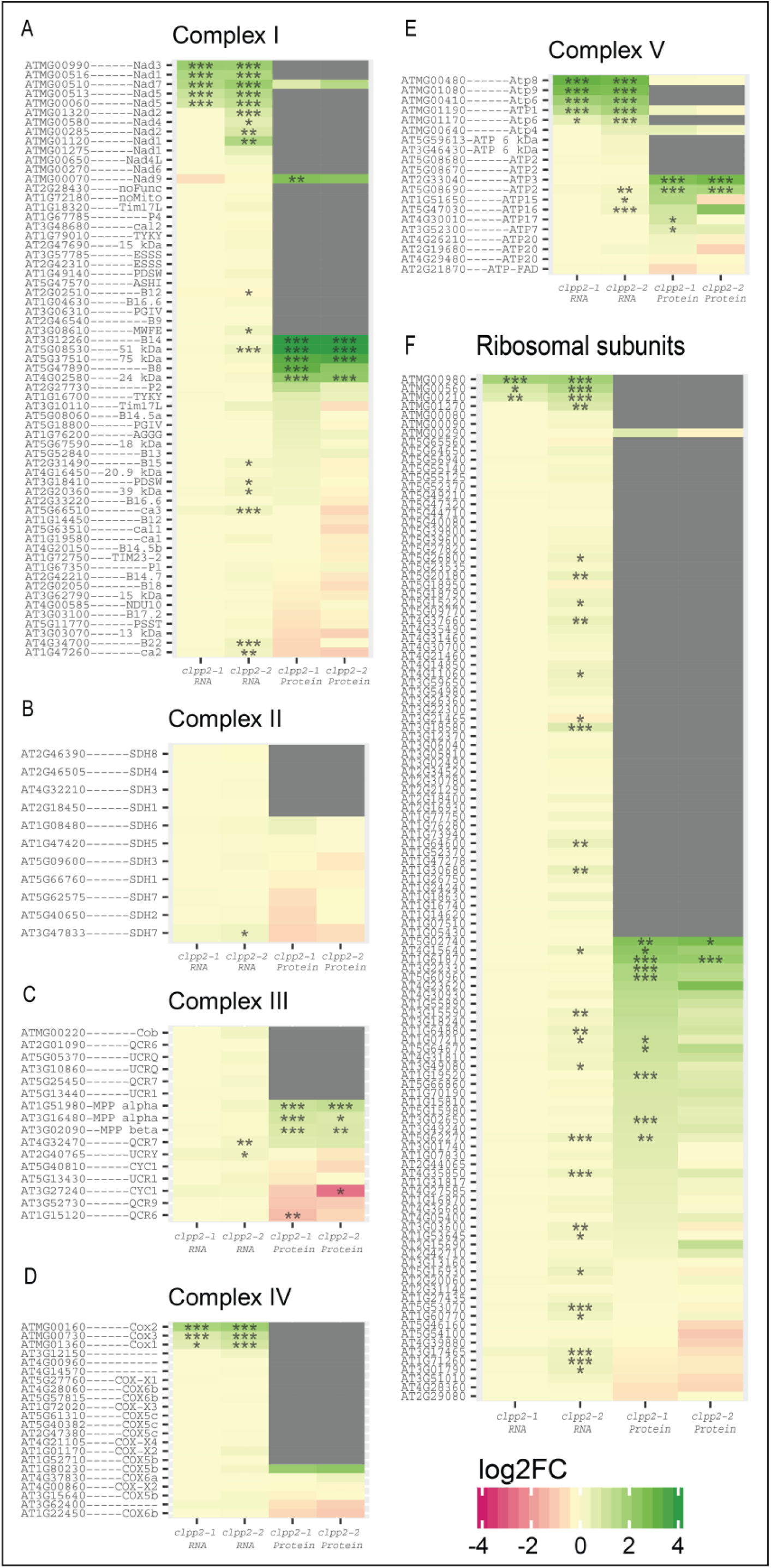
Combination of changes in transcript and protein abundances from OXPHOS and ribosome complexes in *CLPP2* mutants. The selected complexes are: (**A**) Complex I; (**B**) Complex II; (**C**) Complex III; (**D**) Complex IV; (**E**) Complex V and (**F**) the mitochondrial ribosomal complex. The heatmaps show the combination changes in transcript abundance (RNA listed at the bottom) from RNAseq (Figure 2A) and protein abundances (protein listed at the bottom) from quantitative proteomics (Figure 3A). The subunits with accession number and abbreviations from individual complex are listed on the left side. In heatmaps, the colour code represents the log2 fold change of the individual mutants compared to the wild type with green for accumulation and red for depletion in the mutants. Grey colour indicates un-detected proteins. * represent p ≤ 0.05, ** represent p ≤ 0.01, *** represent p ≤ 0.001). (n= 3 for transcript and n = 4 for protein).

Complex III contains only one mitochondrial encoded subunit (Braun and Schmitz, 1992; Meyer et al., 2019) and its expression appeared to be unaffected transcriptionally in both mutants. Changes in protein abundance of MPPα-1, MPPα-2 and MPPβ, were observed but without any changes in transcript level for these nuclear encoded genes (Figure 4C). Complex IV has three mitochondrial subunits, all showed upregulation in transcript abundance in both mutants compared to the WT, and 13 nuclear genes encoded subunits of Complex IV showed no change in transcript or protein abundance (Figure 4D). Complex II of the OXPHOS system contains only nuclear-encoded subunits and appeared to not be affected by a disruption of the mitochondrial CLPP2 protease at the transcript or protein level (Figure 4B). We did not find consistent transcriptional response in both CLPP2 mutants for mitochondria-encoded subunits of the mito-Ribosome (Figure 4F), yet two nuclear-encoded ribosomal large subunits (AT4G30930, AT5G64670) and one nuclear-encoded ribosomal PPR336 (AT1G61870) were increased in protein abundance in both mutants (Figure 4F).

### Loss of CLPP2 impacts protein homeostasis and the assembly of Complex I

Changes in the transcript level and protein abundance of specific Complex I subunits lead us to assess the impact on respiratory chain Complex I in both mutants as an exemplar of a complex affected in *CLPP2* mutants. We firstly analysed Complex I activity in both mutants compared to WT as the rate of de-amino NADH-dependent FeCN reduction but did not find any difference in enzymatic activity (Figure 5A). To determine the abundance of assembled Complex I, we separated mitochondrial complexes using BN-PAGE and stained gels to visualise proteins with Coomassie (Figure 5B). We did not observe any accumulation of Complex I or supercomplex I-III_2_ in either mutant compared with WT (Figure 5B). Complex I in gel activity staining using NADH and NBT as substrate (Yan et al., 2007) showed supercomplex I-III_2_ (band a) and Complex I (band b) had similar in-gel activities in both mutants and WT (Figure 5B), consistent with their protein abundance and total Complex I enzymatic activity. However, we found a residual in-gel activity stain in a lower molecular mass complex (band c) in both mutants but not in WT (Figure 5B). This indicated the existence of a potential subcomplex of Complex I containing a functional FMN cofactor to convert NADH to NAD^+^ and to transfer the free electron onto NBT. To further confirm this hypothesis, we cut the gel band c region from both mutants and WT and subjected them to mass spectrometry analysis. We detected 51 kDa and 24 kDa subunits of Complex I in both mutants but not in WT (Figure 5C), pointing to the existence of a substantial amount of a soluble unassembled N-module subcomplex. We then measured the protein abundance of the three accumulated N-module subunits (24 kDa, 51 kDa and 75 kDa) by targeted multiple reaction monitoring (MRM)-MS in purified mitochondria after fractionation into membrane and soluble fractions by centrifugation. In the membrane fraction a significant 1.5-2-fold accumulation of the 24 kDa and 75 kDa subunit in *clpp2-1* and *clpp2-2* was observed when compared to WT (Figure 5D). However, in the soluble fraction the accumulation was 4-6.5-fold and significant for all 3 subunits (Figure 5D). This indicated that the accumulation of specific Complex I N-module subunits appeared to be an accumulation of an unassembled subcomplex even though the steady-state intact Complex I abundance was not affected in either mutant.

**Figure 5:**
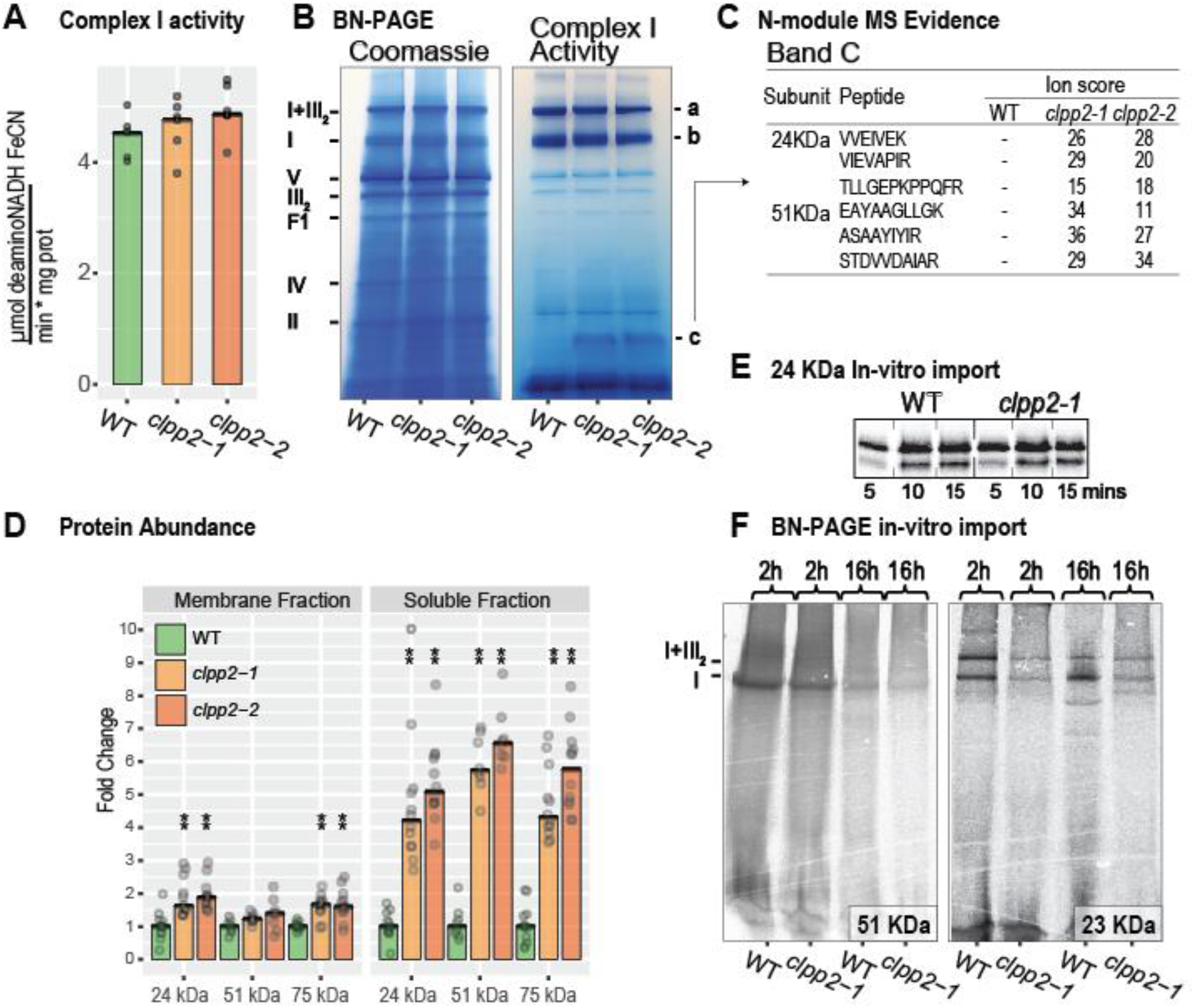
Changes in protein abundance for Complex I N-module subunits and Complex I enzymatic activity and in-vitro import for Complex I assembly in *CLPP2* mutants. (**A**) Complex I enzyme activities in isolated mitochondria from mutants and wild type using de-amino NADH as substrate and FeCN as donor measured using photo-spectrometer assay (n=5). (**B**) BN-PAGE of isolated mitochondria of WT, *clpp2-1* and *clpp2-2*, stained with Coomassie blue was shown in the left and the image of BN-native in gel complex I activity stain (NADH and NBT) was presented in the right. Major OXPHOS components are marked on the gel for size comparison. Bands of interest are marked with a (Supercomplex I-III2), b (Complex I) and c (potential Complex I assembly intermediate). (**C**) Mass spectrometry evidence for N-module proteins (24KDa and 51 KDa) present in band c in the BN-PAGE. (**D**) Protein abundances of three Complex I matrix arm N-module subunits (24 Kda, 51 kDa and 75kDa) in membrane and soluble fractions from isolated mitochondria. Data is shown as fold change in peptide abundance compared to the wild type median. ** represent p ≤ 0.01 (n= 4). (**E**) In-vitro import of radio-labelled 24kDa subunit of Complex I into isolated mitochondria of WT and *clpp2-1* at 5, 10- and 15-minutes incubation time. (**F**) In-vitro import of radio-labelled Complex I 51KDa subunit (left) and 23KDa subunit into isolated mitochondria of WT and *clpp2-1* at 2 hours and 16 hours incubation time. The blue-native (BN) gels were imaged for radioactive intensity. Major OXPHOS components are marked on the gel for size comparison.

We conducted an in-vitro import assay of a radio-labelled 24 kDa subunit to understand the nature of the accumulation of the upregulated Complex I subunits in *clpp2*. There was no apparent difference in the degree of import of the 24 kDa subunit in WT and *clpp2-1* on SDS-PAGE (Figure 5E). Next, we tested for any impairments of Complex I assembly in *clpp2-1* by analysis of radiolabelled imports of the 51 kDa and 23 kDa subunit after 2 and 16 hours using blue native-PAGE (Figure 5F). After 2 hours of 51 kDa import and assembly, we found the emergence of new fully assembled Complex I and supercomplex I-III_2_ in WT and *clpp2-1*, with a higher abundance in WT (Figure 5F). After 16 hours of 51 kDa import and assembly, we observed the same fully assembled Complex I and supercomplex I-III_2_ bands, however they were a lot less abundant (Figure 5F), which could be due to degradation of the imported and assembled protein over time. After two hours of a 23 kDa subunit import we observed similar radiolabelled Complex I and supercomplex I-III_2_ bands in WT and *clpp2-1*, again displaying higher abundances in the wild type (Figure 5F). After 16 hours, we saw the fully assembled Complex I and Supercomplex I-III_2_ bands in wild type and the mutant, but they were relatively similar in abundance to the two hour import and assembly assay, which could be an indication of less degradation than for the 51 kDa protein. We also saw a set of new radiolabelled bands with a lower molecular weight than Complex I after 16 hours of 23 kDa subunit assembly (Figure 5F). The WT showed 2 extra bands, which are both less abundant than the Complex I band and could be assembly or degradation intermediates. We concluded that a Complex I N-module subcomplex cannot be efficiently degraded when CLPP2 is disrupted in plant mitochondria, resulting in its accumulation in the matrix and this in turn may disturb the rate of assembly of new Complex I, but not its final abundance.

### Plants endure the loss of mitochondrial CLPP2 under normal and stress conditions

We evaluated the growth of *clpp2-1* and *clpp2-2* compared to wild type plants under a range of growth conditions to find any phenotypical consequences caused by the loss of the mitochondrial CLPP2. Arabidopsis seedlings, grown for two weeks under long-day control conditions, showed no phenotypic differences between the WT, *clpp2-1* and *clpp2-2* (Figure 6A). In mouse myoblasts, disruption of mitochondrial CLPP is known to lead to an altered mitochondrial morphology (Deepa et al., 2016). Plant roots are a mitochondria rich tissue for energy production without chloroplasts. We employed transmission electron microscopy to investigate the morphology of CLPP2 deficient Arabidopsis mitochondria in root tissues and found no visible differences between the WT, *clpp2-1* and *clpp2-2* (Figure 6B). We grew Arabidopsis seedlings on vertical MS-media agar plates to investigate root elongation under control and various stress conditions. Under control conditions (Figure 6C) and under various stress conditions (salt, osmotic, heat, carbon starvation) (**Supplemental Figure 4**), we did not find significant changes in root elongation between WT and either *clpp2-1* or *clpp2-2*. We also treated plants with inhibitors of Complex I (rotenone), Complex III (antimycin A) and the mitochondrial biogenesis inhibitor (doxycycline), but again no difference in growth was found (**Supplemental Figure 4)**. So far, the only stress condition affecting *clpp2-1* and *clpp2-2* differently when compared with WT was nitrogen starvation after 14 days (Figure 6D), which showed a significantly reduced root growth; about 60% of the WT root length. Therefore, Arabidopsis seedlings can compensate for the loss of CLPP2 under most growth conditions, but under nitrogen starvation scenarios it appears that CLPXP may help to acclimate to stress conditions efficiently.

**Figure 6:**
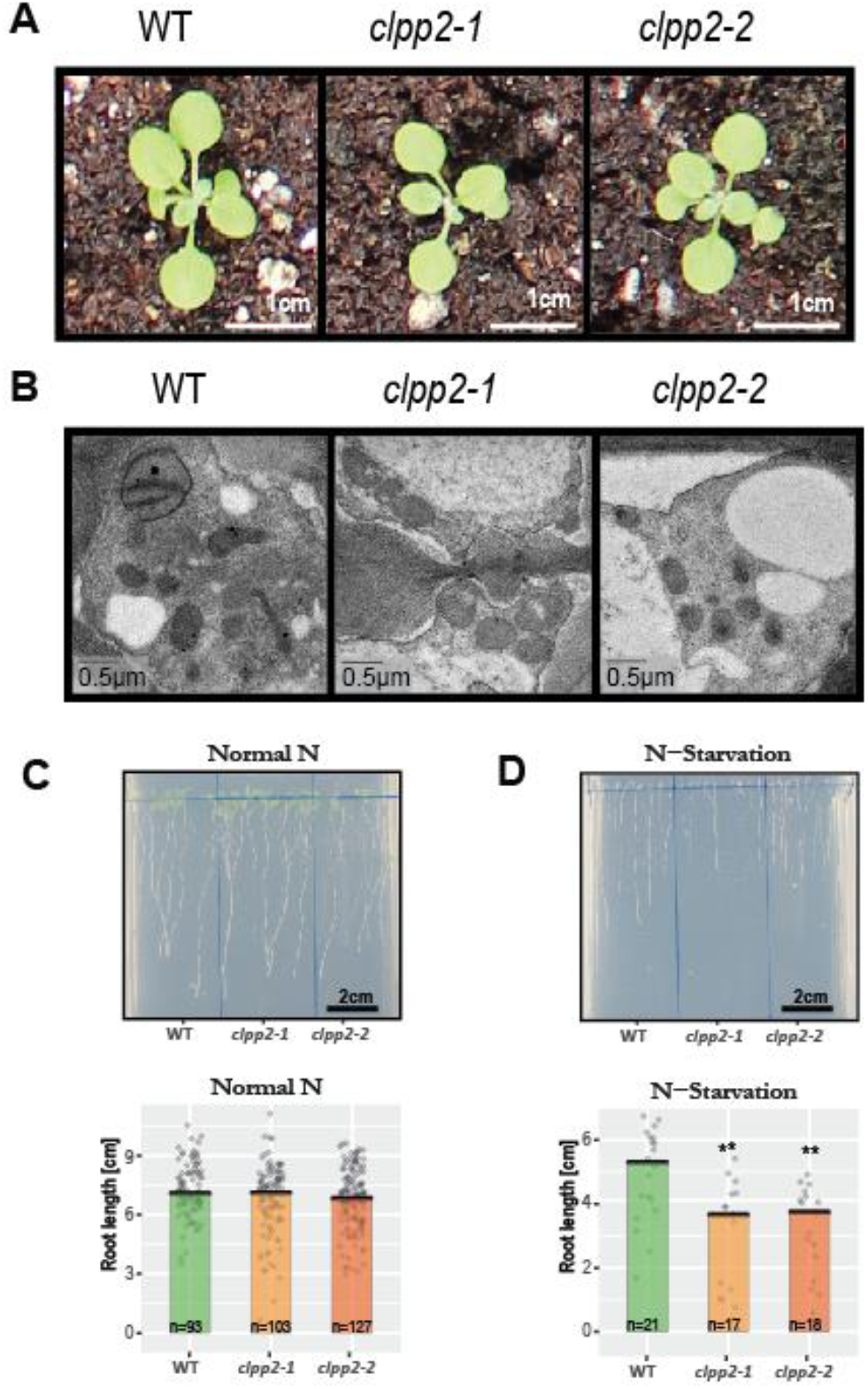
Phenotypes associated with the loss of the mitochondrial CLPP2 under different conditions. (**A**) Arabidopsis thaliana seedlings grown on soil for 15 days under long day conditions of wild type, *clpp2-1* and *clpp2-2*. (**B**) Representative TEM images of dissected roots of 2 weeks old hydroponically grown Arabidopsis thaliana seedlings of wild type, *clpp2-1* and *clpp2-2*. (**C**) Two weeks old *Arabidopsis thaliana* seedlings grown on vertical MS-agar (pH 5.8, 1% sucrose) plates under control long day conditions. Top: Representative plate image. Bottom: Root length measurements, dots represent individual root measurements. (**D**) Two weeks old Arabidopsis thaliana seedlings grown on vertical MS-agar (pH 5.8, 1% sucrose) plates without any nitrogen supply under control long day conditions. Top: Representative plate image. Bottom: Root length measurements, dots represent individual root measurements, ** represent p ≤ 0.01.

## Discussion

CLPP has been well studied in bacteria, some eukaryotes such as human, mice, fungi and in particular in the plant chloroplast where it plays an essential role in chloroplast biogenesis and development (Yu and Houry, 2007; van Wijk, 2015; Bhandari et al., 2018). While it has been long known that a CLPP exists in plant mitochondria (Peltier et al., 2004), there was no information on its function due to lack of genetic resources for its disruption and analysis. In this study, we successfully developed two mutant lines using CRISPR-Cas9 to knockout the Arabidopsis mitochondrial *CLPP2* gene to provide direct insight into its function not only in mitochondrial protein homeostasis but also in coordinated regulation of assembly of plant respiratory complex subunits mixed in their encoding across the mitochondrial and nuclear genomes.

To our surprise, the knockout of mitochondrial *CLPP2* did not cause any phenotypic variation in either CRISPR-Cas9 mutant when compared with WT (Figure 6). In contrast, the plastid-encoded CLPP1 in tobacco (*Nicotiana tabacum*) plays an essential for shoot development (Shikanai et al., 2001). In Arabidopsis, CLPP4 and CLPP5 null mutants were embryo lethal, and null mutants of CLPP3 were seedling lethal in soil. Partial down-regulation of *CLPP4* and *CLPP6* by antisense RNA techniques reduced plant growth and development and resulted in pale-green plants (Sjögren et al., 2006; Zheng et al., 2006; Olinares et al., 2011). Unlike its chloroplast counterpart, the plant mitochondrial CLPP appears to not be essential for plant growth and development. As bacteria lack a proteasome protein degradation system, CLPP and LON proteases contributed to about 80% of their cellular protein degradation (Goldberg et al., 1994). Bacterial CLPP plays an important role in the degradation of proteins involved in nutrient starvation, stationary phase adaptation, heat-stress response, cell-cycle progression, biofilm formation, cell motility, and metabolism (Frees et al., 2003; Gerth et al., 2008). The plant mitochondrial LON protease plays a key role in regulation of mitochondrial protein homeostasis and loss of mitochondrial LON1 causes a severe reduction in plant growth and development (Rigas et al., 2009; Solheim et al., 2012; Li et al., 2016). By contrast, there is no information on the function of Lon protease in chloroplastic protein degradation even though one isoform, LON4, is targeted to the chloroplast (Ostersetzer et al., 2007). The contrasting role of CLPP and LON proteases in chloroplast and plant mitochondria might indicate a shift from Lon protease to CLP protease for the regulation of critical aspects of chloroplast and mitochondrial protein homeostasis, respectively.

Dysfunction of mitochondrial CLPP causes significant phenotypes in other eukaryotes. Loss of mitochondrial CLPP in humans is linked to infertility and sensorineural hearing loss (Gispert et al., 2013; Szczepanowska et al., 2016). Mitochondrial CLPP in humans is required for protein homeostasis (Szczepanowska et al., 2016), which is involved in the degradation and regulation of several enzymes of the electron transport chain and other cellular metabolic pathways such as mitochondrial translation/tRNA modification, mitochondrial transcription, protein folding and proteolysis (Fischer et al., 2013; Cole et al., 2015; Szczepanowska et al., 2016). Loss of mitochondrial CLPP in mice leads to general infertility in both males and females, reduction in physical growth and also sensorineural deafness (Gispert et al., 2013; Szczepanowska et al., 2016). Remarkably, deletion of mitochondrial CLPP leads to improved health and increased life span in the filamentous fungus, *Podospora anserina* (Fischer et al., 2013). In our study, loss of plant mitochondrial CLPP did not have any apparent defect to plant growth and development. Therefore, mitochondrial CLPP has diverse impacts on growth and development of eukaryotic organisms, potentially due to redundant roles in the mitochondrial protease network.

In this study, we observed that loss of CLPP2 only affected the expression of ∼0.1% of nuclear genes (Figure 2A), but nearly 33% of mitochondrial genes (Figure 2A), indicating an important impact of this protease on the landscape of mitochondrial gene expression patterns. Mitochondrial ATP-dependent proteases, such as LON and CLPP, are proposed to serve roles in mitochondrial DNA functions including packaging and stability, replication, transcription and translation (Matsushima and Kaguni, 2012). CLPP may play an indirect role in the regulation of mtDNA replication, transcription and translation (Matsushima and Kaguni, 2012). We observed that loss of CLPP2 caused high expression levels of genes encoding potential RNA/DNA polymerases (ATM00490; ATMG00810) and genes encoding a putative intron maturase (ATMG00520) and a nucleic acid-binging protein (ATMG00560) (Figure 2). In addition, the protein abundance of mitochondrial RNA helicase RH9 (AT3G22310) and RH53 (AT3G22330) (Figure 3 and supplemental Figure 2) increased in both mutants. Similarly, the protein abundance of chloroplast AtRH3 (AT5G26742), which is closely related to mitochondrial RH9 and RH53 (Asakura et al., 2012), was strongly (more than 5-fold) accumulated in in *clpr2-1* and *clpr4-1* (Rudella et al., 2006; Kim et al., 2009; Zybailov et al., 2009). Chloroplast RH3 DEAD Box RNA Helicases in maize and Arabidopsis function in splicing of specific group II introns and affect chloroplast ribosome biogenesis (Asakura et al., 2012). So far, there is no detailed information on the global impact on expression of chloroplast genes caused by loss of chloroplast CLPP, but such studies are likely to be hard to interpret given the severity of the plastid phenotypes of these mutants.

The effect of CLPP2 loss on the mitochondrial accumulation of nuclear-encoded proteins for complexes with mixed subunits derived from both mitochondria and nuclear genomes can be obviously seen for Complex I, Complex V and the mito-ribosome (**Fig4A, 4E and 4F**). This is reminiscent of the previous evidence that a post-translational process is responsible for balancing the availability of nuclear subunits for complex assembly (Sarria et al., 1998; Giege et al., 2005). We have shown through detailed examination of Complex I that this effect is primarily due to an excess subunit availability and not to a block on import, pre-protein processing or accumulation of assembled and functional complexes, indicating that CLPP2 prevents such protein accumulations. It is known that CLPP targets subunits within protein complexes in a range of other organisms, providing a housekeeping mechanism for mixed origin complexes (Flynn et al., 2003; Feng et al., 2013; Fischer et al., 2013; Szczepanowska et al., 2016). Indeed, Arabidopsis orthologs of CLPP substrates identified by trapping approaches in other organisms (including the 24kDa and 75kDa subunits of Complex I) were identified as DEPs in this study (**Supplemental Table 3**).

While Giegé et al. (2005) previously identified the non-stoichiometry of mixed origin subunits phenomenon in a transition from starved to re-fed Arabidopsis callus culture cells, we found here that carbon starvation did not produce a phenotype in *clpp2* plants (Supplemental Figure 4), but transient response to nitrogen starvation did (Fig 6). This implies that non-stoichiometric accumulation of unused subunits is not in itself egregious for plant mitochondrial function, but perhaps it is suboptimal as is shown by the decreased Complex I assembly when N-module accumulates in *clpp2* (Figure 5, Figure 7). The reasons for the specific need for mixed subunit complexes to turnover in this way are unknown. However, there is additional evidence for such maintenance of Complex I. The nuclear-encoded matrix arm subunits of Complex I, including the 75-kDa, 51-kDa, 39-kDa, 18-kDa, and B17.2 subunits, exhibited relatively higher turnover rates than other subunits of Complex I in Arabidopsis (Li et al., 2013). In WT, faster turnover of 51 kDa subunit was observed via *in-vitro* import with decrease radioactive intensity in Complex I after 16 h import in both WT and *clpp2*-1, while radioactive intensity of 23 kDa subunit in Complex I was relatively stable (Figure 5F). Blue native activity assays and targeted proteomic analysis indicated that the N-module subcomplex was highly accumulated in the matrix fraction in both mutants indicating increased stability of this separated part of complex I in *clpp2* plants (Figure 5, Figure 7). In CLPP-deficient mice, reduced protein levels of complex I subunits have also been observed (Deepa et al., 2016; Szczepanowska et al., 2016). Recently, rapid turnover of the N-module in proliferating versus differentiated mammalian cell cultures was found using SILAC, showing higher turnover within blue native PAGE purified intact mitochondrial complexes (Wittig et al., 2018). This suggests that mitochondrial CLPP could have a wider role in this stoichiometric balancing across eukaryotes and assist not only the coordination but also the assembly of mitochondria and nuclear-encoded protein complexes.

**Figure 7:**
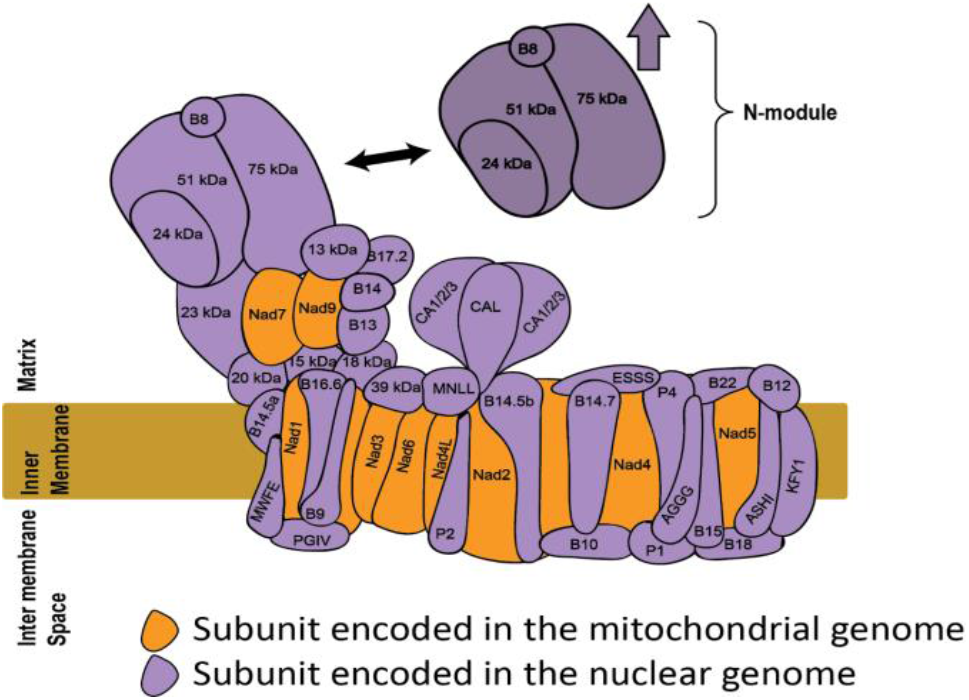
Schematic position of N-module subunits in Complex I and the unassembled N-module subcomplex in the mitochondrial matrix. In both *CLPP2* mutants, the subunits in unassembled N-module subcomplex accumulated at much higher level than those in the assemble holo-complex as indicated by targeted proteomic analysis in soluble and membrane fractions. The presence of highly (∼5-fold) accumulated N-module subunits in matrix, may be related to the reduced assembly of imported 51 KDa subunit and 23 KDa subunit (with physical interaction with N-module subunits) into assembled Complex I and supercomplexI+III (Figure 5E).

## Material and Methods

### CRISPR-Cas9 guided mutagenesis of *clpp2*

The gRNAs were designed for each target at *CLPP2* exon1 using CRISPRdirect (Naito et al., 2015) and further analysed with E-CRISPR (Heigwer et al., 2014). The cassettes containing two gRNAs for each mutant were ordered as gBlocks and were inserted into the pHEE2E backbone, provided by Dr Qi-Jun Chen (Wang et al., 2015), by restriction-ligation using BsaI (NEB) (**Supplemental Figure 1**). All plants were *Arabidopsis thaliana* plants (ecotype Columbia-0), grown on soil as described below, and were transformed by Agrobacterium-mediated DNA insertion using the floral dipping procedure (Clough and Bent, 1998). Plants transformed with the CRISPR-Cas9 construct were selected by resistance to hygromycin (15 µg/ml). Genomic DNA was extracted from leaf tissue through a protocol adapted from (Edwards et al., 1991). Primers F1: ATGAGGGGTCTCGTTTCCG and R1: GCCGCCAGGGGAATTGAGA were used for PCR genotyping. Homozygous T1 plants were determined by the presence of a single trace including desired mutations after Sanger sequencing. T2 plants were selected for the absence of the CRISPR-Cas9 T-DNA by PCR. T2 and T3 seedlings were used for subsequent experiments.

## Plant growth

Arabidopsis (*Arabidopsis thaliana*) seeds (Columbia-0: WT, CRISPR-Cas9 KO mutants: *clpp2-1* and *clpp2-2*) were sown on a perlite:vermiculate:compost (Seedling Substrate Plus+,Bord Na Móna, Ireland) soil mix (1:1:3) and covered with a transparent hood. After vernalization for 3 days at 4 °C in the dark, plants were grown under control conditions: long-day photoperiod (16h light: of 150 µmol m^−2^ s^−1^ light intensity, 8 h dark),70% relative humidity, 22°C during the day and 18°C at night. For selection of transformed plants, seeds were spread on Murashige and Skoog Gamborg B5 plates (0.8% (w/v) agar, 1% (w/v) sucrose, 0.05%(w/v) MES, pH 5.8-KOH), supplemented with 15 μg/ml hygromycin B, and placed in the dark at 4°C for 3 days. Then plates were transferred into long day conditions as described above. For root length measurements, seeds were surface sterilized and placed on Murashige and Skoog Gamborg B5 plates as described above in vertical orientation. For root stress treatments, Arabidopsis seedlings were grown on plates as described above, with addition or removal of specific growth conditions: rotenone (addition of 5µM rotenone), antimycin A (addition of 50µM antimycin A), doxycycline (addition of 1mg/l doxycycline), salt stress (addition of 100 mM NaCl), osmotic stress (addition of 175 mM mannitol), carbon starvation (no sucrose supplied), heat (seedlings on plates grown at 30°C day/night).

For mitochondria isolation and RNA extraction, Arabidopsis (*Arabidopsis thaliana*) seeds were surface sterilised and dispensed into one-half strength Murashige and Skoog Gamborg B5 liquid media, within enclosed sterilized 100ml polypropylene containers. The containers were seated on a rotating shaker in the long-day conditions as described above and seedlings were harvested two weeks after germination.

### RNAseq analysis

RNA was extracted from three biological replicates per genotype using the SIGMA Spectrum™ Plant Total RNA Kit according to the instructions. The RNA quality and quantity were assessed using a ThermoFisher nanodrop ND-1000 spectrophotometer (ThermoFisher, USA). RNAseq library preparation and 50bp single-end read sequencing on BGISEQ-500 was performed by BGI Group, China. RNAseq data were received from BGI as fastq file-format. Following quality control with FastQC (https://www.bioinformatics.babraham.ac.uk/projects/fastqc/), on average, 42 million reads were obtained per sample, which were mapped to Arabidopsis reference genome Ensembl, v34 and quantified with Salmon (v0.14.0) (Patro et al., 2017). Differential analysis was performed with the R package DESEQ2 (Love et al., 2014).and transcripts with a log2 fold change between mutant and WT (log2FC) exceeding ±0.4 and an adjusted p-value less than or equal to 0.05 were considered differentially expressed.

RNA sequencing data has been deposited to the Gene Expression Omnibus (Barrett et al., 2012) with the accession number GSE141942.

### Mitochondria isolation, protein purification and trypsin digestion

The two week old hydroponically grown Arabidopsis seedlings were used for mitochondria isolation (Lee et al., 2008). The isolated mitochondria were quantified using Bradford method (Bradford, 1976) and the aliquots were stored at −80°C until further analysis.

For MRMs used to confirm the CLPP2 loss at the protein level, 200 µg mitochondria were precipitated with 9x volumes ice-cold acetone for 24h at −20°C and was centrifuged at 20,000 x g for 20 minutes at 4°C. The pellets were collected for further analysis. For quantitative untargeted MS and MRM-MS analysis of different mitochondrial fractions, 200 µg mitochondria were lysed using 3x freeze-thaw cycles (20min at −20°C and then 20 mins at 4°C) and centrifuged at 20,000 x g for 20 minutes at 4°C. The supernatant (soluble fraction) and pellet (membrane fraction) were collected and then precipitated with 9x volumes cold acetone for 24h at −20°C. The pellets from soluble and membrane fractions were collected for further targeted MS analysis using MRM. For quantitative untargeted MS and MRM-MS experiments, samples were alkylated and trypsin digested as follows:

The above acetone precipitated pellets were resuspended with 100µl solution containing 50mM ammonium bicarbonate, 10 mM dithioreithol (pH 8.0) and incubated at 60°C for 30 mins. Samples were cooled to room temperature and alkylated with 100µl 50mM ammonium bicarbonate, 25mM Iodoacetamide for 30 minutes. Samples were trypsin digested by adding digestion solution (1:50 (w/w, trypsin/protein) trypsin, 50mM ammonium bicarbonate, 2% (v/v) acetonitrile, 1.2mM CaCl_2_, 0.1M guanidine GuHCl, pH 8.0) and incubated at 37°C for 16 hours in a thermomix at 1000 rpm. Digested samples were desalted and concentrated using C18 macroSpin columns (The Nest Group, USA) following the manufacturer instructions and then eluted with 100µL solution (80% acetonitrile, 0.1% formic acid). Elutes were dried under vacuum, resuspended in 2% (v/v) acetonitrile, 0.1% (v/v) formic acid to a final concentration of ∼1µg µL^-1^ protein. Finally, samples were filtered through Ultrafree-MC Centrifugal Filter (0.22µm, PVDF) following the instructions of the manufacturer (MilliporeSigma, USA).

### Quantitative untargeted mass spectrometry

Samples were analysed by LCMS on a Thermo orbitrap fusion mass spectrometer using data dependent acquisition. Analysis consisted of direct injection onto a self-packed 150 mm x 75 µm Dr Maisch Reprosil-Pur 120 C18-AQ 1.9 µm column. Water/acetonitrile gradients with 0.1% formic acid were formed by an Ultimate U3000 nano pump running at 250 nL min^-1^ from 2-27% acetonitrile over 30 minutes.

Thermo raw files were database searched and quantified using MaxQuant (v1.6.7.0) (Cox and Mann, 2008) and analysed using the R package DEP (Zhang et al., 2018). Based on PCA analysis sample one WT sample and one clpp2-2 sample were flagged as outliers (high overall variance compared to all other samples) and removed.

Quantitative untargeted mass spectrometry have been deposited to the ProteomeXchange Consortium via the PRIDE (Perez-Riverol et al., 2019) partner repository with the dataset identifier PXD016746. Reviewer account details: Username: Password: vRUlkOnv.

### Targeted MRM mass spectrometry

MRM-MS was conducted as described in (James et al., 2019), in brief: 10 μL of each sample were loaded onto an Agilent AdvanceBio Peptide Map column (2.1 × 250 mm, 2.7 μm particle size, P.N. 651750-902), using an Agilent 1290 Infinity II LC system. The column was heated to 60°C. Peptides were eluted over a 15 minute gradient (0-1 min 3% [v/v] acetonitrile 0.1% [v/v] formic acid to 8% [v/v] acetonitrile 0.1% [v/v] formic acid; 1-15 min 8% [v/v] acetonitrile 0.1% [v/v] formic acid to 45% [v/v] acetonitrile 0.1% [v/v] formic acid; 15-15.5 min 45% [v/v] acetonitrile 0.1% [v/v] formic acid to 100% [v/v] acetonitrile 0.1% [v/v] formic acid; 15.5-16 min 100% [v/v] acetonitrile 0.1% [v/v] formic acid to 3% [v/v] acetonitrile 0.1% [v/v] formic acid; 16-30 minutes 3% [v/v] acetonitrile 0.1% [v/v] formic acid) into the Agilent 6495 Triple Quadrupole MS for detection. Peptide transitions used for multiple reaction monitoring are given in **Supplemental Table 1.**

### Protein labelling with iTRAQ reagents, ChaFRADIC amd LC-MS/MS analysis of N-terminal peptides

In total, 12 samples corresponding to four biological replicates of each genotype i.e. WT, *clpp2-1* and *clpp2-1* were used and divided into two experiments. Briefly: isolated mitochondria pellets (∼100 µg of protein) of each genotype and their respective biological replicates were first solubilized with 10 µL of 10% SDS containing complete (mini protease inhibitor cocktail) and samples were diluted to 100 µL with 50 mM NH_4_HCO_3_ buffer (pH 7.8). Reduction and subsequent alkylation steps were carried out with 10 mM dithiothreitol and incubation at 56°C for 30 min; followed by alkylation of free thiol groups using 30 mM iodoacetamide and incubation at room temperature for 30 min in dark. Next, each sample was diluted 10-fold with ice-cold ethanol in 1:10 ratio, vortexed and stored at −40°C for 60 min followed by centrifugation in a pre-cooled (4°C) centrifuge at 18,000 g for 30 min. Next, the supernatant was discarded and to each protein pellet 100 µL of ice-cold acetone were added, briefly vortexed and centrifuged as above. The supernatant was discarded and the protein pellets were dried under a laminar flow hood. Labelling of proteins with iTRAQ reagents, enrichment of the N-terminal peptides based on the ChaFRADIC method, LC-MS/MS and data analysis were performed as previously described (Venne et al., 2013; Venne et al., 2015).

Mass spectrometry proteomics data have been deposited to the ProteomeXchange Consortium via the PRIDE (Perez-Riverol et al., 2019) partner repository with the dataset identifier PXD016263. Reviewer account details: Username: Password: 2y7IvNFO.

### Sequence logo analysis

Sequence logos of N-terminal peptides detected by ChaFRADIC MS were produced using the R package ‘ggseqlogo’. Sequence logos are created from frequency data, while the ChaFRADIC MS results finally yield intensity data. The intensities were normalized and transformed into frequency data. RH9 (AT3g22310) and RH53 (AT3g22330) were identified as outliers (contributed >50x sequences) and removed for this calculation. Transformation and normalization were performed using the following rules. Normalization: Raw peptide intensity / median all intensities of the peptide. Transformation: Normalized intensity <= 0.1 : Frequency = 1, Normalized intensity <= 0.2 : Frequency = 2, and then using 0.1 increments for 1 Frequency to a maximum Frequency of 25. Sequences were then multiplied based on their frequencies and the sequence logo was calculated.

### Complex I enzymatic activity assay

Complex I enzymatic assays were carried out as described previously (Huang et al., 2015). Deamino-NADH:Q reductase (Complex I) specific activity was measured by the following reaction. The reaction solution contained Tris-HCl, 50 mM pH 7.2; NaCl, 50 mM; FeCN, 1 mM; deamino-NADH, 0.2 mM. The decrease of absorbance at 420 nm at 25°C was recorded after adding 10 µg mitochondrial protein in 1 ml reaction solution. The extinction coefficient of FeCN at 420 nm is 1.03 mM^-1^ cm^-1^.

### Blue-Native gel separation and Complex I in gel activity staining

Three hundred µg mitochondrial proteins were dissolved with 5mg digitonin/mg protein and then separated with 4.5-16% gradient BN-PAGE gel (Schertl and Braun, 2015). A part of gel was stained with Coomassie blue. For Complex I in gel activity assay, the gel was washed three times for 5 mins with distilled water and incubated in the reaction medium (0.14 mM NADH, 1.22 mM NBT, 0.1 M Tris-HCl, pH7.4). When the dark blue stain was strong enough, the reaction was stopped by transferring gel to 40% methanol/10% acetic acid (v/v) (Schertl and Braun, 2015).

### In-gel spot identification

Gel spots (∼1mm^3^) of interest were excised from the blue-native gel and destained twice in appropriate volumes of 50%(v/v) methanol. 50%(v/v) 50mM ammonium bicarbonate, 0.1%(v/v) formic acid for 30min at 1000rpm and twice dehydrated in appropriate volumes of 50%(v/v) acetonitrile / 50% (v/v) 50mM ammonium bicarbonate, 0.1%(v/v) formic acid for 1minute and 30 seconds in 100% acetonitrile. Samples were then reduced, alkylated, dehydrated, digested and cleaned up as described above in appropriate volumes and identified using an Agilent 6550 Q-TOF as described in (Nelson et al., 2014).

### Protein import assays

[^35^S]-Methionine-labelled proteins were translated using the rabbit reticulocyte TNT in vitro transcription translation kit (Promega) using the 24 kDa (AT4G02580) subunit cloned into pDEST14 as previously described (Ivanova et al., 2019). In vitro protein import assays were carried out as described previously into 30 µg of freshly isolated mitochondria (Duncan et al., 2015). BN-PAGE import assays were carried out using [^35^S]-methionine-labelled proteins as previously described (Duncan et al., 2015). In brief: radiolabelled protein was incubated with 30 µg of freshly isolated mitochondria for 2h or 16 h, the mitochondria was pelleted by centrifugation and processed for BN-PAGE analysis (Eubel et al., 2005), except that commercially available precast native-PAGE gels were used (Novex Tris-glycine, Thermofisher). The 23 kDa (AT1G79010) and the 51 kDa (AT5G08530) subunits of Complex I were cloned using Gateway technology into pDONR201 and subsequently recombined into pDEST14 for [^35^S]-methionine-labelled protein translation as described above.

### Transmission electron microscopy

The root tissues of two week old hydroponic seedlings were used and the analyses were performed in Australian National University’s Electron Microscope laboratory referring to the method described previously (Hyman and Jarvis, 2011). The dissected root tissues were fixed overnight in 2.5% (v/v) glutaraldehyde/ 2% (v/v) paraformaldehyde in 0.1 M sodium phosphate buffer (pH 7.2), and then post-fixed with 1% (w/v) OsO4/1.5% (w/v) potassium ferricyanide. The tissues were then dehydrated in gradual ethanol and propylene oxide, embedded in Spurr’s resin and cured for 24 hours at 70 degree. Ultrathin sections of approximately 70nm thickness were cut using a Leica UC7 ultramicrotome, collected onto copper mesh grids, and stained with 1% uranyl acetate and Reynolds’ lead citrate. The images taken under a Hitachi7100 transmission electron microscope at 75kV. Images were acquired using a Gatan Orius CCD camera.

## Acknowledgement

J.P. was supported by UWA International Fee Scholarships; S.H., A.P and E.C. were supported by an Australian Research Council Discovery Grant (DP140101580). A.H.M, and R.L. and J.C were supported by Australian Research Council (CE140100008). R.L. was supported by a Sylvia and Charles Viertel Senior Medical Research Fellowship and Howard Hughes Medical Institute International Research Scholarship. S.H. and M.W.M. were supported by Australian Research Council Future Fellowships (FT130101338 and FT130100112, respectively). A.P. was supported by Australian Research Council Discovery Early Career Research Award (DE120102913). L.K., S.W. and A.S. acknowledge the support by the Ministerium für Kultur und Wissenschaft des Landes Nordrhein-Westfalen, the Regierende Bürgermeister von Berlin - inkl. Wissenschaft und Forschung, and the Bundesministerium für Bildung und Forschung. J.L. was supported by the National Collaborative Research Infrastructure Strategy (NCRIS).

## Supporting Documents

**Supplemental Figure 1:** Cloning history of the plasmid to create the CRISPR-Cas9 construct used to create stable *CLPP2* knockout lines.

**Supplemental Figure 2:** Changes in protein abundances in soluble fraction of isolated mitochondria in *CLPP2* mutants compared with WT.

**Supplemental Figure 3:** Sequence logos of N-terminal peptides obtained from ChaFRADIC MS of purified mitochondrial proteins from WT, *clpp2-1* and *clpp2-2*.

**Supplemental Figure 4:** Arabidopsis seedlings of wild type, *clpp2-1* and *clpp2-2* grown on MS-agar plates (pH 5.8, 1% sucrose) under stress conditions or additional chemicals.

**Supplemental Table 1:** Peptide transition list are used for multiple reaction monitoring (MRM) mass spectrometry analysis.

**Supplemental Table 2:** Changes in protein abundances in membrane and soluble fraction of isolated mitochondria in *CLPP2* mutants compared with WT.

**Supplemental Table 3:** List of subunits in mitochondrial complexes as substrates for CLPP targets through trapping approaches in other organisms.

